# Metal-induced energy transfer (MIET) imaging of cell surface engineering with multivalent DNA nanobrushes

**DOI:** 10.1101/2023.07.05.547790

**Authors:** Dong-Xia Wang, Bo Liu, Gui-Mei Han, Qing-Nan Li, De-Ming Kong, Jörg Enderlein, Tao Chen

## Abstract

The spacing between cells has a significant impact on cell-cell interactions, which are critical to the fate and function of both individual cells and multicellular organisms. However, accurately measuring the distance between cell membranes and the variations between different membranes has proven to be a challenging task. In this study, we employ metal-induced energy transfer (MIET) imaging/spectroscopy to determine and track the inter-membrane distance and variations with nanometer precision. We have developed a DNA-based molecular adhesive called the DNA nanobrush, which serves as a cellular adhesive for connecting plasma membranes of different cells. By manipulating the number of base pairs within the DNA nanobrush, we can modify various aspects of cell-cell interactions, such as adhesive directionality, distance, and forces. We demonstrate that such nanometer-level changes can be detected with the MIET imaging/spectroscopy. Moreover, we successfully employ MIET to measure distance variations between a cellular plasma membrane and a model membrane. This experiment does not only showcase the effectiveness of MIET as a powerful tool for accurately quantifying cell-cell interactions, but does also validate the potential of DNA nanobrushes as cellular adhesives. This innovative method holds significant implications for advancing the study of multicellular interactions.

## Introduction

Cell-cell interaction is a vital physiological process in multicellular organisms.^1^ Accurate control of cell-cell interaction presents a way to study and manipulate various cellular processes, thus benefiting the development of cell-based theranostics and tissue engineering. ^2,3^ Cell surface engineering strategies hold great potential in regulating cell-cell interactions by modifying the surface with various functional materials, such as proteins, nucleic acids, nanoparticles, or polymers.^4–7^

While cell surface-engineered materials or surface-engineering strategies are experiencing thriving development,^8,9^ a crucial aspect that has been largely overlooked is the characterization of their ability to regulate the spacing between membranes. This oversight can be attributed to the limited availability of characterization tools specifically designed for this purpose. However, understanding and characterizing the intermembrane spacing^10–12^ and cell fusion. ^13,14^ Therefore, it is imperative to develop suitable tools and techniques to investigate nanomaterial interactions at nanometer distances with cellular membranes, as this knowledge is essential for comprehending cell membrane surface engineering processes.

Various methods and techniques, such as cryogenic transmission electron microscopy (cryo-TEM),^15,16^ neutron reflectometry (NR),^17^ or superresolution fluorescence microscopy^18^ have been employed to quantify the distance between different membranes. However, those methods and techniques are limited in their ability to observe dynamic changes during adhesion processes. Although reflection interference contrast microscopy (RICM) and total internal reflection fluorescence microscopy (TIRFM) have the potential to measure membrane dynamics,^19–21^ their application to nucleated cells is risky due to the presence of complex components inside the cell and proteins on the membrane. These factors can lead to ill-defined variations in refractivity, making the measurements unreliable.

Recently, our group has developed a new method called metal-induced energy transfer (MIET) to precisely determine the axial position of a fluorescent single molecule above a metal film.^22,23^ The principle of MIET is based on the energy transfer of the excited state energy of an excited fluorophore to surface plasmons in the metal film. This energy transfer is extremely distance-dependent and leads to a distance-dependent modulation of fluorescence lifetime and intensity.^24^ This mechanism is similar to Förster resonance energy transfer (FRET), where an emitter (the donor) transfers its excited state energy to a nearby molecule (the acceptor), provided that the emission spectrum of the donor overlaps with the absorption spectrum of the acceptor. Due to the broad absorption spectra of metals, the energy transfer from a fluorescent molecule to the metal film takes place with high efficiency across the full visible spectrum. Thus, any dye in the visible spectral range will be affected by MIET, and its measured fluorescence lifetime can be converted into a distance of the emitter from the metal surface. MIET has been used for investigating various systems, from whole cells to organelles, and to determine the axial position of individual molecules with a precision of ca. 3 nm.^25,26^

Here, we use MIET imaging/spectroscopy to precisely measure the intermembrane distance in DNA-nanostructured modulated membrane systems with nanometer-scale accuracy (Scheme 1). To achieve this, we have developed a DNA-based adhesive called a DNA nanobrush. One of the key advantages of this DNA adhesive is its versatility in design,which allows for the manipulation of valence states to modify intercellular forces between cell membranes. Additionally, it provides the ability to adjust the number of base pairs on the brush backbone and tentacles, thereby enabling controlled regulation of adhesive directionality and distance. We demonstrate that MIET can effectively and accurately measure nanometer-sized changes in the distance between cell membranes decorated with the DNA nanobrushes. Furthermore, we apply MIET to monitor the adhesion process between cellular membranes induced by the DNA nanobrush. Our results do not only confirm the potential of MIET as a powerful tool for studying intercellular interactions but do also highlight the potential of our DNA nanobrush as a molecular glue for cellular assembly.

**Scheme 1:**
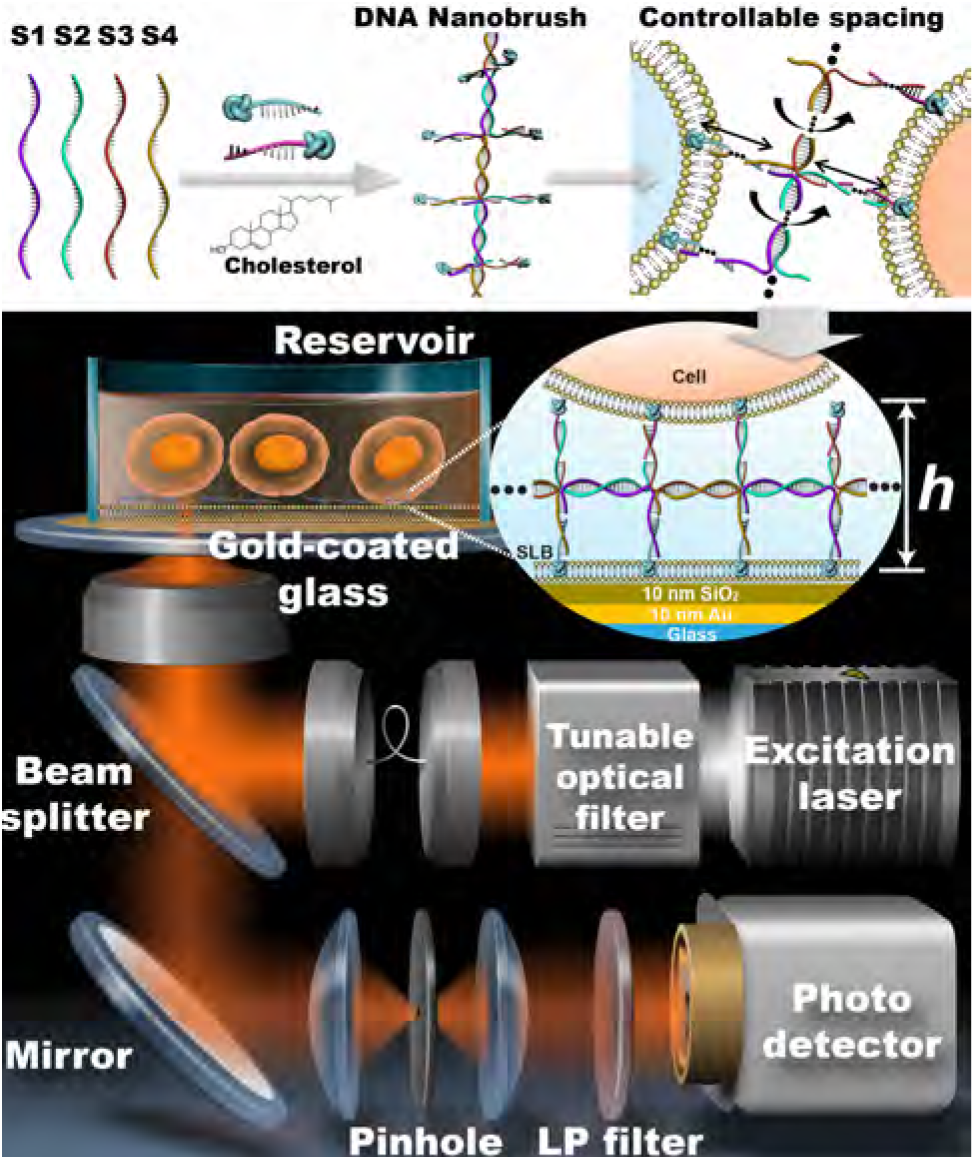
Multivalent DNA nanobrush engineered on a cell membrane surface for precisely quantifying intercellular interactions via metal-induced energy transfer.

## Results and discussion

### Synthesis and characterization of multivalent DNA nanobrushes

A series of DNA nanobrushes were designed and synthesized for cell membrane surface engineering (see SI experimental section and Table S1). The backbone units of these DNA nanobrushes were composed of four short DNA strands (S1, S2, S3, and S4) linked in series to form a linear core (Figure 1A). The number of base pairs (bps) in the linear backbone is designed to be 21 bp (b1); 22 bp (b2) and 25 bp (b3), respectively. Theoretically, each base pair contributes about 0.34 nm of length and about 34.3° of twist to the growing helix,^27^ resulting in a helical twist of 10.4 base pairs/turn (bp/turn) for B-form DNA.^28^ Thus, the base pair numbers that one uses determines the arrangement and direction of the functional strands: gradually from a planar (21 bps, b1) to a twisted brush (b2 and b3, Figure 1B and Figure S1). From the nanobrush backbone, numerous side arms (tentacles) can extend that can hybridize with cholesterol-labeled complementary strands, thereby introducing cholesterol functional groups that can link to a membrane. By using different single-stranded sequences on the side arms, it is possible to artificially control the number and position of introduced cholesterol groups. Here, we employ two different single-stranded sequences on the side arms, each capable of hybridizing with its respective cholesterol-labeled complementary strand. When hybridized with one of these complementary strands, cholesterol is added to one half of the side arms (b1-1chol). When hybridized with both complementary strands simultaneously, cholesterol is added to all side arms (b1-2chol). Compared to other rigid DNA structures such as DNA origami or amphiphilic DNA probes, the DNA nanobrush offers more flexibility, which is advantageous for interaction with highly flexible membranes, even when they exhibit high curvature and dynamics. ^29,30^

**Figure 1:**
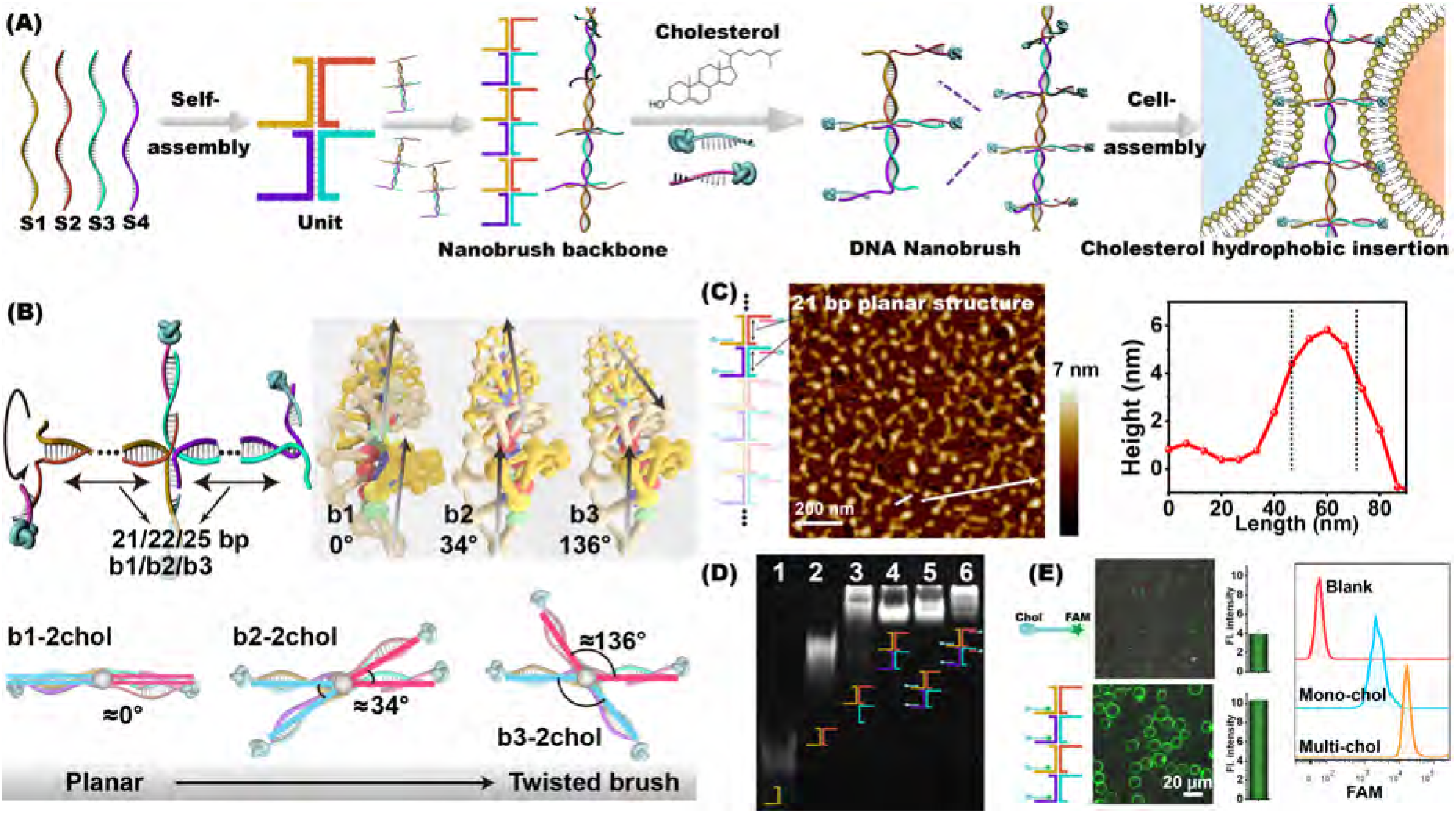
Synthesis and characterization of the DNA nanobrushes. (A) Schematic illustration of cell surface engineering with a DNA nanobrush. (B) Structural illustration of DNA nanobrushes (b1, b2, b3) with different twist angles due to changing backbone. (C) AFM images of a DNA nanobrush (b1-2chol). The right panel shows a linear cross-section of the height profile of a nanobrush, displaying dimensions of approximately 6 nm in height and 30 nm in diameter. (D) PAGE assay of different DNA samples. From lanes 1 to 6: S1; S1+S2; S1+S2+S3; S1+S2+S3+S4 (brush backbone, b1); b1-1chol; b1-2chol. (E) Membrane anchoring capacity of monovalent-cholesterol and multivalent-cholesterol DNA nanobrushes. The middle panels show fluorescence intensity images. The right panel shows a flow cytometry analysis of monovalent- and multivalent cholesterol. More than 100 cells per group were measured and then analyzed with ImageJ, and for each group, three independent measurements were performed.

We first synthesized various functional DNA nanobrushes through a self-assembly process using a linear backbone composed of four short DNA strands, and arms modified with cholesterol functional groups to link membranes. First, the structure of the DNA nanobrush was evaluated with atomic force microscopy (AFM). Using b1-2chol DNA nanobrush as an example, as shown in Figure 1C, b1-2chol shows a flexible linear structure with a length of ∼100 nm, demonstrating the successful assembly of the nanobrush. The height of the brush is only ∼6 nm and the width is ∼30 nm, which suggests a planar structure for the b1 DNA nanobrush. Polyacrylamide gel electrophoresis (PAGE) demonstrates the successful self-assembly of a nanobrush by rapid programmable annealing (Figure 1D). By sequentially adding DNA from S1 to S4, the gel migration of the DNA mixture slows down gradually (Lines 1-4). After adding the functional side arm strands (lanes 5 and 6), all gel bands with delayed migration correspond to samples with successful assembly. Next, we evaluated the anchoring ability of the nanobrush to cell membranes. We attached FAM fluorophores to cholesterol-labeled DNA strands. CCRF-CEM, a T lymphoblastoid cell line, was used as a model cell line. With multivalent hydrophobic vertices, DNA nanobrushes (b1-1chol) showed strong membrane-anchoring ability without becoming internalized. Over 90% of the probe was retained on the membrane even after incubation in 10% FBS-containing medium for one hour. However, the monovalent-cholesterol strand (single-strand DNA labeled cholesterol) rapidly dissociated from the cell membrane after 30 minutes (Figure S2). As evaluated from the fluorescence intensity, multivalent-cholesterol DNA nanobrushes are almost 2.7 times larger in thickness than the monovalent-cholesterol ones (Figure 1E). This was further confirmed with flow cytometry. These results demonstrate that multivalent cholesterol can stably anchor nanostructures on the membrane surface, providing a stable anchoring method for subsequent MIET measurements.

### Nanobrush for cell surface engineering

We utilized our nanobrush to guide and program cell-cell binding versatility, using CCRF-CEM cells and Ramos cells (human B lymphoma cells) as test samples. We optimized bonding conditions using the planar structure (b1-2chol) with a nanobrush backbone of 21 bp (Figure S3). Confocal microscopy imaging and flow cytometry (Figure 2A) revealed no assembly between the two cells without the addition of the nanobrush. However, in the presence of a multivalent cholesterol-modified DNA nanobrush, the assembly efficiency reached 66.4%, indicating that our DNA nanobrush does significantly improve cellular adhesion efficiency between different cell types.

**Figure 2:**
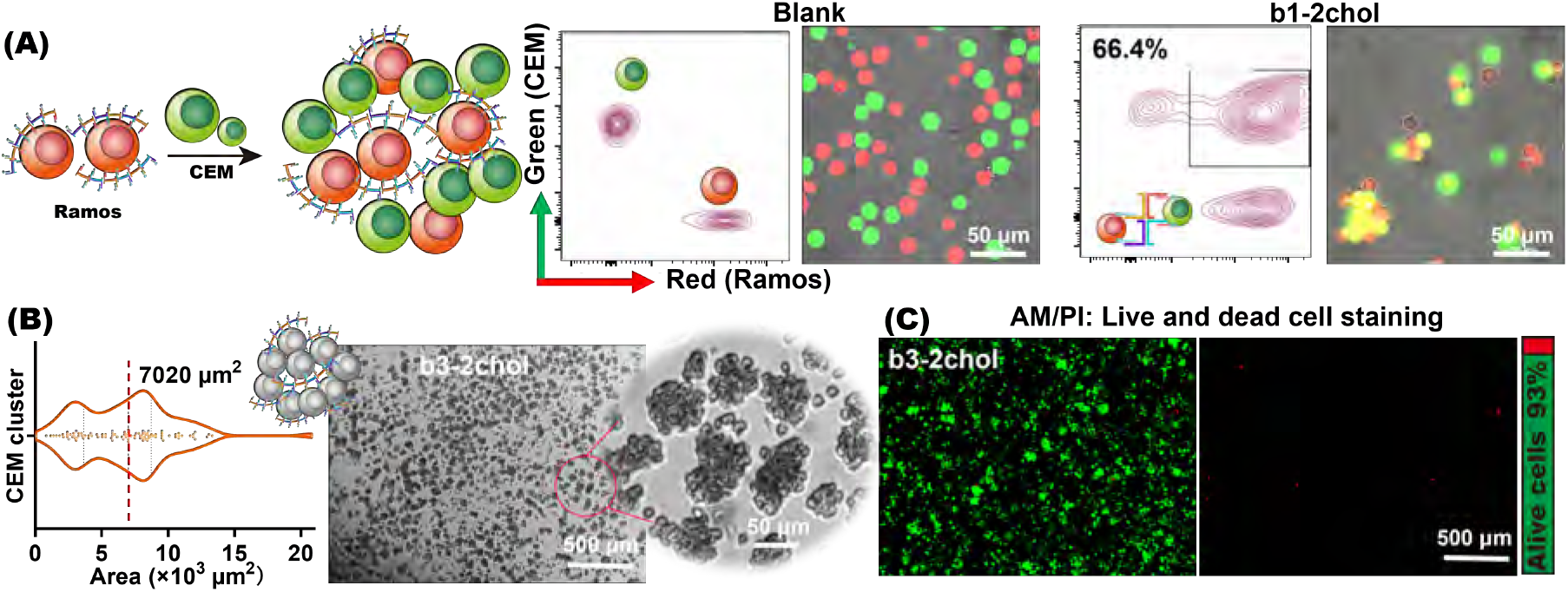
Nanobrush for cell surface engineering. (A) DNA nanobrush (b1-2chol) induced heterotypic cell aggregation. Assembly characterization was performed using confocal laser scanning microscopy (CLSM) and flow cytometry, respectively. Green: CCRF-CEM cells; Red: Ramos cells. (B) Aggregates of CCRF-CEM cells assembled with b3-2chol. The cross-sectional area distribution of homogeneous CEM cell spheroids after assembly for 24 h. The area of 100 cell clusters was measured using ImageJ. Characterization was performed using bright field cell microscopy. (C) Live-dead staining image of CEM cell spheroids after 36 h. Alive and dead cells were stained with Calcein-AM (green) and PI (red), respectively. The bar on the right shows the quantified optical density of the fluorescence images.

Next, we investigated the potential of our DNA nanobrush for adhering homogeneous cells to form stable cell clusters and differentiate into microtissue over suitable incubation times. Here, we used a nanobrush (b3-2chol) with a backbone base of 25 bp. As shown in Figure 2B, after 24 h incubation, the b3-2chol nanobrush regulated CEM cells to form a larger cluster. The median cross-sectional area was determined to be about 7020 μm^2^ by counting 100 clusters, corresponding to 50 - 150 cells per cluster. Live-dead cell staining experiments demonstrated that most cells within the spheroids were still alive after 36 h incubation (Figure 2C).

### DNA nanobrush-regulated inter-membrane distance determined by MIET imaging/spectroscopy

Having established DNA nanobrushes as an effective and reliable nanoglue for plasma membranes that works both between homogeneous cells and heterogeneous cells, we applied MIET imaging/spectroscopy to determine the inter-membrane distance, which is modulated by our DNA nanobrushes. A biomimetic membrane system was designed to model cell adhesion between supported lipid bilayers (SLBs, prepared using DOPC (1,2-dioleoyl-sn-glycerol-3-phosphocholine)), and fluorescently labeled giant unilamellar vesicles (GUVs, prepared with DOPC and 0.1% DPPE-Atto655 (1,2-Bis(diphenylphosphino)ethane)).

To validate the high spatial resolution of MIET imaging/spectroscopy, a series of DNA nanobrushes with different arm lengths were placed between the SLBs and GUVs. SLBs were prepared on a MIET substrate (10 nm gold film is sandwiched between a coverslip and a 10 nm silica layer) via vesicle fusion. Then, DNA nanobrushes were added and incubated for 30 min to form DNA nanobrush layers on the SLBs (Figure S4). After washing out the unbounded DNA nanobrushes, fluorescently labeled GUVs were added to the chamber and incubated for another 30 min. The gold film-coated substrate served for inducing a distance dependent fluorescence lifetime. Fluorescence images and lifetimes of the fluorescently labeled GUVs were taken with a confocal microscope, which was equipped with time-correlated single-photon counting (TCSPC) for fluorescence lifetime measurements.^31^

We first scanned the sample and found that the proximal membranes of almost all GUVs adhered to the SLBs via the DNA nanobrushes. In contrast, GUVs without any DNA brushes did not exhibit adhesion events (Figure S5). To precisely determine the inter-membrane distance, we scanned individual GUVs to accumulate signal for fluorescence decay fitting. We constructed TCSPC curves for each pixel and fitted these curves with a multi-exponential decay model, giving us a mean fluorescence lifetime for each pixel (Figure S6).^32^ These lifetime images were then converted to membrane height images above the silica surface using a MIET calibration curve (Figure 3A). A MIET calibration curve was calculated based on a semi-classical electrodynamics model of the near-field coupling between the fluorophore and the substrate, taking into account parameters such as the refractive index of the buffer, the thickness of the metal and silica films, and the quantum yield and emission spectrum of the dye molecules. For the system used here, all the optical parameters for both the dye DPPE-Atto655 (free-space lifetime = 2.60, quantum yield = 0.36) and the Au/SiO_2_ substrate have been published before (more details for calibration curve calculation are given in the Supporting Information). ^24,33^ Figure 3B shows three calculated MIET curves for DPPE-Atto655 for three different dye orientations with respect to the 10 nm Au film with 10 nm spacer. Based on the lifetime images, we observed that the proximal membrane shows a uniform height across the SLB surface with no height fluctuations. Consistent with our previous findings,^23,34,35^ in the case of a flat planar membrane, we assume a dye orientation parallel to the membrane, with the dipole axis of Atto655 parallel to the surface, which is important for the lifetime-height conversion.

**Figure 3:**
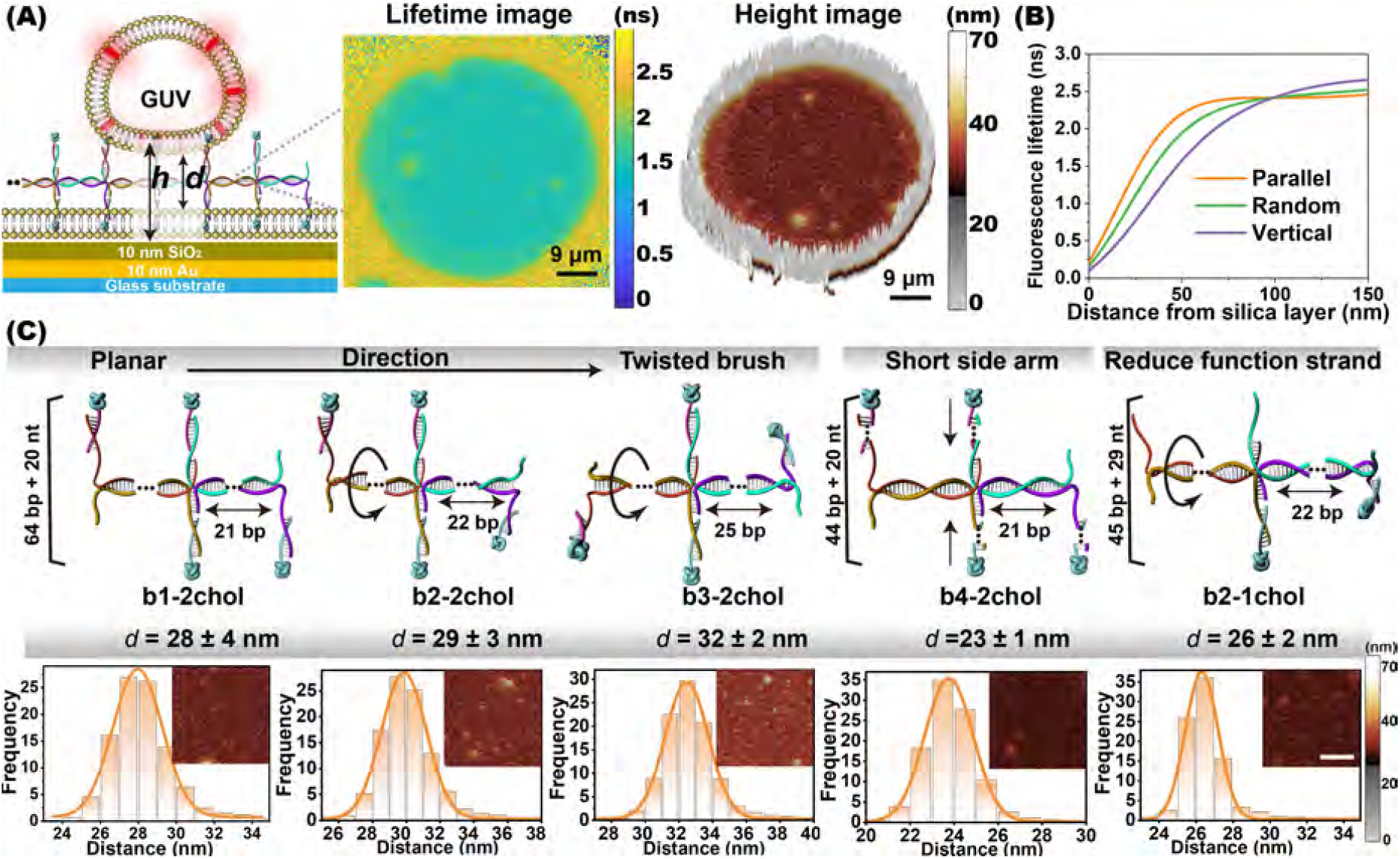
Metal-induced energy transfer (MIET) microscopy to visualize the inter-membrane distance regulated by DNA nanobrushes. (A) Schematic illustration of MIET imaging/spectroscopy to measure the distance between a GUV and SLB membranes modulated by DNA nanobrushes. *h* is the height from the center of the proximal membrane of one GUV to the SiO_2_ surface. *d* is the distance between the two membranes. Middle panel: fluorescence lifetime image obtained from calculating the mean lifetime with a maximum likelihood estimation algorithm for each pixel. Right panel: height map *d* of the proximal membrane of a GUV. (B) Calculated dependence of fluorescence lifetime on axial distance from silica surface. Curves were calculated for a dipole emitting at a wavelength of 680 nm and for three different dipole orientations with respect to the interface (vertical, horizontal, random orientation). The MIET substrate was fabricated by depositing 10 nm of gold and 10 nm of SiO_2_ on a cover slide. (C) DNA nanobrushes regulate the average distance *d* between a GUV and an SLB. Orange curves show Gaussian fits of the distance distributions. More than 10 GUVs per group were measured, and for each group three independent measurements were performed. Inset: corresponding height map. Scale bar is 9 μm.

Figure 3A shows a height image measured from the b1-2chol/GUV system. The image displays a uniform attachment of the proximal membrane of the GUVs to the nanobrush. The controllable and uniformly distributed cholesterol-functioned strands on the nanobrush backbone facilitate an even distribution of cholesterol over the membrane surface, preventing the formation of self-aggregating clusters. The bright white color of the GUV’s circumference reflects the substantial distance of its membrane there from the gold surface, which exceeds the measurement range of MIET. For statistical analysis, we quantified the height values or distances of all pixels in the central region and calculated the average values (Figure 3C). The height (*h*) values of the central area were found to be uniform within a range of 34 nm to 39 nm. Taking the hydration layer of the SLBs (∼2 nm) and the thickness of the lipid bilayer (∼4 nm) into account, the inter-membrane distance (*d*_*m*−*m*_) was derived by subtracting 8 nm from the height (*h*).

To further evaluate the sensitivity of MIET imaging/spectroscopy for detecting subtle changes in DNA nanobrush structure, we repeated experiments for different nanostructure with varying backbone and side arms. We first varied the number of backbone bases (b1; b2; b3) to regulate the arrangement direction of the functional strands, gradually from a planar to a twisted brush. Interestingly, even though the nanobrushes (b1; b2; b3) had the same arm lengths (64 bp + 20 nt), the resulting inter-membrane distance gradually increases from the planar to the twisted conformation (from 28 ± 4 nm to 32 ± 2 nm). This can be attributed to the increasing rigidity of the nanostructure in the rotating conformation, reducing the tilt of the DNA structure between two membranes. Secondly, we modified the nanobrush by shortening the side arms (b4) for the planar structures. As expected, a reduction of 20 bp in the side arm bases corresponded to a decrease in the distance by approximately 5 nm. Theoretically, 20 bps correspond to 6.8 nm in length, and the observed 5 nm reduction suggests an inclination of the planar structure. ^36^ In addition, we also changed the number of cholesterol functional strands (b2-1chol). When the number of cholesterol functional strands decreased, the distance decreased accordingly (26 ± 2 nm). Reducing one side arm results in a decrease of 10 nt. Additionally, the functional side arm transitions from a double-stranded structure to a single-stranded structure, thereby enhancing the flexibility of the nanobrush. This increased flexibility allows the nanobrush to flatten and adhere to the membrane surface, resulting in a shorter inter-membrane distance. These results demonstrate that MIET can accurately discern the nanoscale changes in cell membrane spacing induced by modifications of DNA nanobrushes.

### Real-time observation of cell surface engineering by MIET imaging/spectroscopy

After demonstrating that MIET imaging/spectroscopy can measure the inter-membrane distance between model membranes, we utilized MIET next to monitor distance changes during DNA-nanobrush mediated plasma membrane adhesion of a single cell to a SLB. We replaced GUVs with NIH-3T3 cells (mouse embryonic fibroblast cells) and selected the nanobrush (b1-2chol) as the nanoglue (Figure 4B). The cell’s plasma membrane was labeled by fusing fluorescently labeled fusogenic liposomes with the membrane.^37^ After fluorescent labeling and washing, the cells were added to the DNA-nanobrush modified SLB, which was supported by an Au/SiO_2_ substrate. Once the cell settled onto the SLB, we started to continuously scanning the sample at a scanning rate of 0.4 s/frame over an area of 20 μm *×* 20 μm. For extracting the dynamics, we obtained TCSPC data for each pixel by frame binning. Then, the fluorescence lifetime values of these pixels were determined with an mono-exponential decay model using a maximum likelihood algorithm. Finally, we convert the measured fluorescence lifetime values of each pixel into height values using the MIET curve. Supporting movie S1 shows the height variations over time of one NIH-3T3 cell during adhesion mediated by the DNA nanobrush at a frame rate of 4 s/frame. As the apical cell membrane is at least 500 nm away from the substrate, only dye molecules within the basal membrane were efficiently excited and detected.

**Figure 4:**
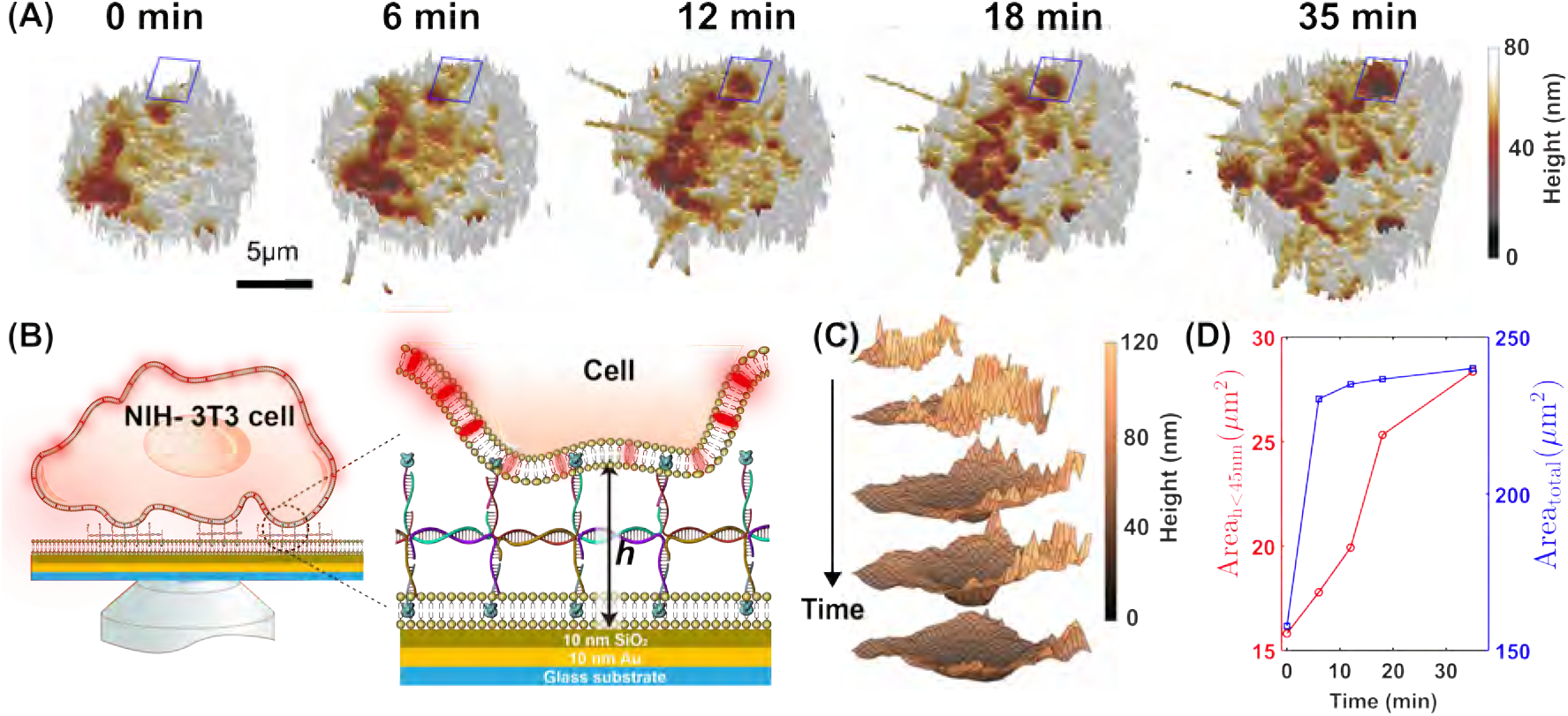
Measurement of DNA-nanobrush regulated binding of a NIH-3T3 cell to an SLB with MIET imaging/spectroscopy. (A) Reconstruction of the 3D height (*h*) maps of the proximal membrane of a NIH-3T3 cell mediated by DNA nanostructures. For each image, the photons are accumulated for 40 s. (B) Schematic illustration of the measurement of a NIH-3T3 cell above SLB with DNA nanobrushes. (C) The enlarged areas are marked in panel A. (D) The statistical analysis of the areas for the contact zones at different times. The red line is the area for the membrane having a height smaller than 45 nm and the blue line is the total area for the whole contact zone.

For a more precise determination of the height values, we constructed height maps by accumulating one frame over 40 s, so that at least 500 photons per pixel contribute to the lifetime calculation. As shown in Figure 4A, the DNA nanobrushes lead to a gradual adhesion of the cell’s plasma membrane to the SLBs. At the beginning of the process, the majority of the membranes are situated at higher elevations with a mean height of ∼70 nm, where only a small fraction with a height of ∼40 nm undergoes adhesion. Over time, both the total area of the contact zone and the area of the adhesion zone (height smaller than 45 nm) increased (Figure 4A, D). A representative enlarged region as shown in Figure 4C provides a detailed illustration of the adhesion process: the plasma membrane adheres to the SLB, and then the adhesion area expands. Adhesion is probably triggered by membrane fluctuation, resulting in a high probability that membrane patches encounter the cholesterol groups of the DNA nanobrush. After the plasma membrane adheres to the DNA nanobrush, the lowered height of the plasma membrane induces subsequent adhesion of neighboring membrane areas, leading to an extension of the adhesion zone. Finally, we checked that the adhesion of the plasma membrane to the SLB is induced by the DNA nanobrush by conducting control experiments without nanobrush. In the absence of DNA nanobrushes, the 3T3 cells did not adhere to the SLB even after 60 min of incubation (Figure S7).

## CONCLUSION

In summary, we have demonstrated the capability of MIET imaging/spectroscopy for monitoring membrane interface changes mediated by DNA nanobrushes. By designing DNA nanobrushes with varying orientation, distance, valence, and flexibility, we used MIET to elucidate subtle changes of membrane spacing as regulated by these DNA nanostructures. Importantly, MIET enables the observation of the adhesion as well as the adhesion dynamics between two membranes of live cells. Additionally, MIET is straightforward to implement and requires neither any hardware modification of a FLIM system nor the preparation of complex sample substrates. Coating glass cover slides with a thin metal film is the only pre-requisite of the technique. The successful utilization of MIET for measuring inter-membrane spacing opens up new avenues for exploring cell surface engineering on the nanoscale.

### Experimental

#### Preparation of DNA nanobrush

All DNA nanobrushes were synthesized through a “one-pot” process. Briefly, six oligonucleotides (Table S1) with identical molar concentrations were mixed in 20 mM Tris-HCl buffer (pH 8.0) containing 50 mM MgCl_2_. The mixtures were heated at 95 °C for 10 min and then incubated on ice for 10 min. The as-prepared nanobrushes were stored at 4 °C for further use.

#### MIET Instrument and preparation of MIET substrate

FLIM measurements were performed using a home-built confocal microscope equipped with a high numerical aperture objective lens (Apo N, 100X oil, 1.49 NA, Olympus Europe, Hamburg, Germany). A pulsed linearly polarized laser (640 nm) with a tunable filter (AOTFnC 400.650-TN, Pegasus Optik GmbH, Wallenhorst, Germany) was used for fluorescence excitation. The light was directed toward the objective through a non-polarizing beam splitter, and back-scattered excitation light was blocked with long-pass filters (FF01-692/40, Semrock). The emission light was focused onto the active area of an avalanche photodiode (PDM Series, MicroPhoton Devices) through a pinhole (100 μm), and the detection times of recorded single photons were determined using a multichannel picosecond event timer (HydraHarp 400, PicoQuant GmbH, Berlin, Germany). A fast Galvo scanner (FLIMbee, Picoquant) was used for imaging scanning. The MIET substrate comprised of a multilayer structure consisting of consecutive layers of 2 nm Ti, 10 nm Au, 1 nm Ti, and 10 nm SiO_2_ on a glass coverslip. This layers were deposited by evaporation using an electron beam source (Univex 350, Leybold) under high vacuum conditions (∼10^−6^ mbar). During the vapor deposition process, the film thickness was monitored using an oscillating quartz unit and then verified by atomic force microscopy.

#### Vesicles and SLB preparation

SUVs were prepared with an extrusion method. Briefly, 100 μL of 10 mg/mL DOPC lipids in chloroform were dried in a vacuum for 1 h at 30 °C to remove the residual solvent. Then, 500 μL of PBS buffer (pH = 7.4) was added and and the solution was shaken for 1 h at 30 °C. The solution was then extruded for 15 cycles through a polycarbonate filter (Whatman) with 50 nm pore diameter. The resulting vesicle solutions were used within 3 days while stored at 4 °C before use. GUVs were fabricated by electroformation ^23^ in a custom-built Teflon chamber with two stainless steel electrodes. Briefly, 100 μL of a chloroform solution containing DOPC (10 mg/mL) and 0.1 M Atto655-DPPE was deposited onto two electrodes, followed by evaporation for 3 h under vacuum at 30 °C. The chamber was filled with 500 μL of 300 mM sucrose solution, after which an alternating electric current of 15 Hz frequency and a peak-to-peak voltage of 1.6 V was applied for 3 h, followed by a lower frequency voltage of 8 Hz for another 30 min. Formed GUVs were collected by rinsing the electrode surface with 500 μL of PBS solution. DOPC SLBs were formed via vesicle fusion. Before placing a SUV solution onto a MIET substrate, the substrate was activated with the plasma of a plasma cleaner (Harrick Plasma, New York, United States) at low intensity for 30 s. Then, a droplet of SUV solution was placed on the substrate and incubated for 1 h to ensure the formation of a uniform bilayer with minimal defects. This was followed by copious washing with buffer.

#### Cell membrane staining

For cell membrane staining, liposomes were prepared by mixing DOPE/DOTAP/Atto655-DPPE lipids in chloroform in a weight ratio of 1:1:0.1. The chloroform was then evaporated under vacuum for 0.5 hours, and the lipids were dispersed in 20 mM HEPES buffer to obtain a final concentration of 2 mg/mL. The solution was vortexed for approximately 2 min to produce multilamellar liposomes. After homogenization in an ultrasonic bath for 20 min, liposomes ready for cell membrane staining were obtained. For cell membrane staining experiments, 5 μL of liposome stock solution was diluted 100 times with appropriate cell culture medium and gently shaken for 1 min at room temperature. Then, ∼10^5^ NIH-3T3 cells were incubated in 500 μL of fusogenic liposomes solution (pH 7.4) for 20 min at 37 °C. Subsequently, the cells were washed twice with 500 μL of 1 × PBS buffer and then suspended in 1 mL of fresh medium for future use.

## Acknowledgement

T.C. and J.E. acknowledge financial support by the European Research Council (ERC) for financial support via project “smMIET” (grant agreement no. 884488) under the European Union’s Horizon 2020 research and innovation program. J.E. acknowledges financial support by the DFG through Germany’s Excellence Strategy EXC 2067/1-390729940. D.K. and D.W. acknowledge financial support by the National Natural Science Foundation of China (No. 22074068). D.W. acknowledges the scholarship from China Scholarship Council.

## Supporting Information Available

The supporting information is available free of charge. Reagents, Characterization, Control experiments, Working principle of MIET imaging, Table for DNA sequences, Schematic diagram of DNA nanobrush, Supporting figures S1-S7.

## TOC Graphic

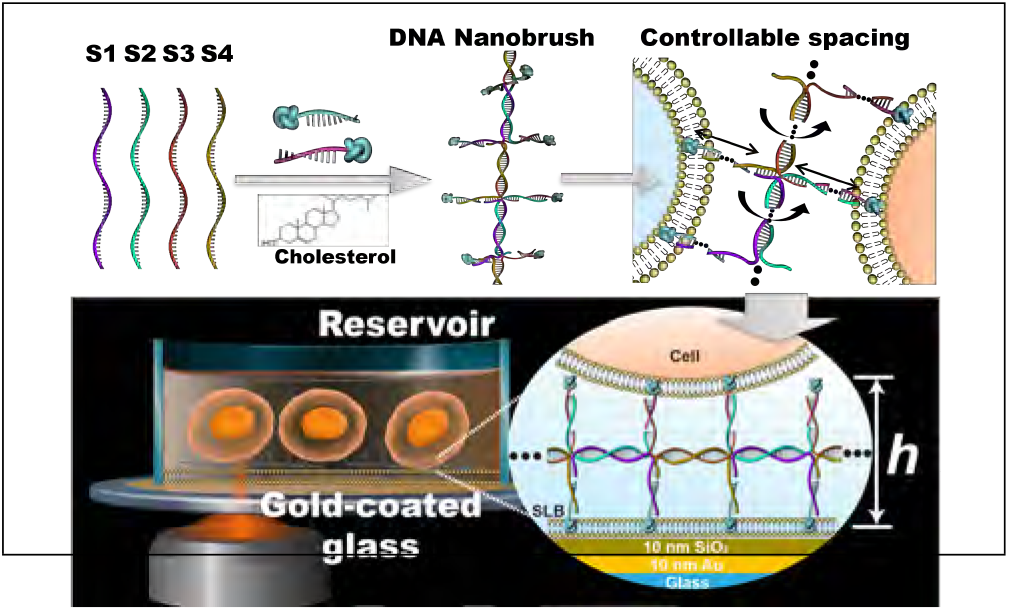

## Notes

### Competing Interest Statement

The authors have declared no competing interest.

