## Supplementary Information for "Metal-induced energy transfer (MIET) imaging of cell surface engineering with multivalent DNA nanobrushes"

### Table of Contents

**Experimental section………………………………………………………………S-1**

**Schematic diagram of DNA nanobrush……………………………………………S-8**

**Synthesis and characterization of the DNA nanobrush.......................................S-11**

**Optimization of two kinds of cells assembled by nanobrush..............................S-12**

**Scheme of MIET experimental set-up…………………………………………S-13**

**MIET measurements of DNA nanobrush-assembled membranes......................S-14**

**MIET microscopy to visualize binding process using NIH-3T3 cells..................S-16**

**References…………………………………………………………………………S-17**

### Video supporting information

**Video S1:** Height variations of one NIH-3T3 cell during the adhesion process mediated by the DNA nanobrush.

### 1. Experimental section

#### 1.1. Materials and reagents

All DNA oligonucleotides (Table S1) were synthesized and purified by Sangon Biotech. Co. Ltd. (Shanghai, China). CM-DiO (green), CM-DiI (red), CellTracker Green CMFDA (green), and CytoTraceTM Red Fluorescent Probe (red) were purchased from Yeasen (Shanghai, China). Calcein-AM and propidium iodide (PI) were obtained from Meilunbio (Dalian, China). Fetal bovine serum (FBS), Roswell Park Memorial Institute (RPMI) 1640, and Dulbecco's modified Eagle's medium (DMEM) were purchased from Gibco (Grand Island, NY, USA). 1,2-dioleoyl-sn-glycero-3-phosphatidylcholine (DOPC), 1,2-dioleoyl-3-trimethylammonium-propane, chloride salt (DOTAP), and atto655 labled 1,2-Bis(diphenylphosphino)ethan (atto655-DPPE) were purchased from Sigma-Aldich (Germany). All other reagents were analytical grade and used directly without further purification. Ultrapure water (resistance ⩾ 18.2 MΩ·cm) was used throughout the experiments.

#### 1.2. Apparatus

The cell fluorescence images were captured using a confocal laser scanning microscope (Nikon A1R) equipped with 4×, 20×, 60× and 100× objective lenses. For MIET measurements, all measurements were carried out with a homebuilt confocal microscope equipped with a multichannel picosecond event timer (HydraHarp 400, PicoQuant GmbH) allowing for fluorescence lifetime imaging. Atomic force microscopy (AFM) characterization was carried out using a Bruker Dimension Icon (USA). Flow cytometry results were recorded on a FACS Calibur (BD, USA).

**Table S1.** Sequences of the oligonucleotides used in this work.

| Oligonucleotide | | Sequences (5' to 3') |
|---|---|---|
| **Nanobrush backbone 21 bases (b1)** | S1-21 | GAAAGAAACAACCCTTGCGCATTCGAGTCTCCACATTTAGTTCTCATCCTTTCCATTCGAGTCTCC |
| | S2-21 | TATTGTTTCTTGGTAGTCTTTAGCTGCAGTGTGAAAGGATGAGAACTAAATGTTTAGCTGCAGTGT |
| | S3-21 | GAAAGAAACAACCCTTGCGGGAGACTCGAATGTAGTATCCCACATTCTCTTTCGGAGACTCGAATG |
| | S4-21 | TATTGTTTCTTGGTAGTCTACACTGCAGCTAAGAAAGAGAATGTGGGATACTAACACTGCAGCTAA |
| **Nanobrush backbone 22 bases (b2)** | S1-22 | GAAAGAAACAACCCTTGCGCATTCGAGTCTCCCACATTTAGTTCTCATCCTTTCCATTCGAGTCTCC |
| | S2-22 | TATTGTTTCTTGGTAGTCTTTAGCTGCAGTGTGAAAGGATGAGAACTAAATGTGTTAGCTGCAGTGT |
| | S3-22 | GAAAGAAACAACCCTTGCGGGAGACTCGAATGTTAGTATCCCACATTCTCTTTCGGAGACTCGAATG |
| | S4-22 | TATTGTTTCTTGGTAGTCTACACTGCAGCTAAGAAAGAGAATGTGGGATACTAAACACTGCAGCTAA |
| **Nanobrush backbone 25 bases (b3)** | S1-25 | GAAAGAAACAACCCTTGCGCATTCGAGTCTCCCACATTTAGTGCGTCTCATCCTTTCCATTCGAGTCTCC |
| | S2-25 | TATTGTTTCTTGGTAGTCTTTAGCTGCAGTGTGAAAGGATGAGACGCACTAAATGTGTTAGCTGCAGTGT |
| | S3-25 | GAAAGAAACAACCCTTGCGGGAGACTCGAATGTTAGTATCCCATGCCATTCTCTTTCGGAGACTCGAATG |
| | S4-25 | TATTGTTTCTTGGTAGTCTACACTGCAGCTAAGAAAGAGAATGGCATGGGATACTAAACACTGCAGCTAA |
| **Nanobrush backbone 21 bases short side arm (b4)** | s-S1-21 | ACCCTTGCGCATTCGAGTCTCCACATTTAGTTCTCATCCTTTCCATTCGAGTCTCC |
| | s-S2-21 | TGGTAGTCTTTAGCTGCAGTGTGAAAGGATGAGAACTAAATGTTTAGCTGCAGTGT |
| | s-S3-21 | ACCCTTGCGGGAGACTCGAATGTAGTATCCCACATTCTCTTTCGGAGACTCGAATG |
| | s-S4-21 | TGGTAGTCTACACTGCAGCTAAGAAAGAGAATGTGGGATACTAACACTGCAGCTAA |
| **Functional strand** | chol-1 | CGCAAGGGTTGTTTCTTTCTTTATCTAAC-chol |
| | chol-1-FAM | FAM-CGCAAGGGTTGTTTCTTTCTTTATCTAAC-chol |
| | s-chol-1 | CGCAAGGGTTTTATCTAAC-chol |
| | chol-2 | AGACTACCAAGAAACAATATTTATCTAAC-chol |
| | s-chol-2 | AGACTACCATTTATCTAAC-chol |

The green and yellow parts represent the backbone of the nanobrush, with S1 and S2 being complementary, and S3 and S4 being complementary as well. The side arms are depicted in red and blue parts, respectively.

##### 1.3. Preparation and characterization of DNA Nanobrush

All DNA nanobrushes were synthesized through a “one-pot” process. Briefly, six oligonucleotides (Table S1) with identical molar concentrations were mixed in 20 mM Tris-HCl buffer (pH 8.0) containing 50 mM $MgCl_2$. The mixtures were heated at 95 °C for 10 min and then incubated on ice for 10 min. The as-prepared nanobrushes were stored at 4 °C for further use. The successful synthesis of the above DNA nanobrushes was validated through polyacrylamide gel electrophoresis (PAGE) gel. 2 μL 6 × Super GelRed Prestain Loading Buffer (US EVERBRIGHT INC.) was added to the above samples and sufficiently mixed. The samples were analyzed by 10% PAGE in 1 × TAE buffer at a constant voltage of 120 V for 50 min. The gel was visualized using a Gel Image System (Amersham Biosciences).

The successful synthesis of nanobrush was also analyzed using atomic force microscopy (AFM). To do this, 1 μM DNA nanobrush samples in a total volume of 50 μL were deposited onto a fresh mica surface. The samples were allowed to adsorb for 15 minutes. After that, 30 μL of ultrapure water was added to the sample and allowed to stay on the surface of the mica sheet for 1 minute. The sample was then absorbed with filter paper, and the washing step was repeated more than 10 times. Finally, the sample was dried with compressed air. The samples were imaged by atomic force microscopy in air scan mode.

##### 1.4. Multivalent and monovalent nanobrush binding CEM cells

For monovalent cholesterol-binding CEM cells, $1\times10^5$ CEM cells were seeded 24 h in advance. After centrifugation to remove the complete medium, 500 μL of 1 × PBS buffer and 100 nM cholesterol-FAM were added and the cells were incubated for 15 minutes at 37 °C. The cells were washed twice with 500 μL of 1 × PBS buffer and then suspended in 1 mL of PBS buffer. To achieve multivalent cholesterol-binding CEM cells, the cholesterol-FAM was simply replaced with nanobrush-FAM. Fluorescence microscopy (Nikon A1R) was used for cell imaging, with the laser excitation wavelength set at 490 nm for FAM. The cell images were observed using a 100× oil objective lens.

##### 1.5. Planar nanobrush for two kinds of cell assembly

Human acute lymphoblastic leukemia CCRF-CEM (abbreviated as CEM) cells and human Burkitt lymphoma Ramos cells were cultured in RPMI 1640 medium (GIBCO). The medium was supplemented with 10% fetal bovine serum (FBS) and 1% antibiotics (penicillin-streptomycin-amphotericin B) and the cells were incubated at 37 °C in a $CO_2$ incubator. $1\times10^6$ Ramos cells were mixed in 5 mL of 1640 medium, puted into a culture flask, and cultured overnight for 12 hours. Then, 1 mL of Ramos cells were taken and centrifuged at 1200 rpm for 3 min. The cells were resuspended with 100 μL PBS and stained with CytoTraceTM Red Fluorescent Probe (red). The cells were then adjusted to $1\times10^5$ for future use. Similarly, for CEM cells, $1\times10^5$ cells were stained with CellTracker Green CMFDA (green) and used for subsequent experiments.

When the two kinds of cells were assembled, differently structured nanobrush (b1) at a final concentration of 500 nM were added to $1\times10^5$ Ramos cells and incubated for 30 min at 37 °C with a metal bath shaking at 300 rpm. The nanobrush was first anchored to the Ramos cell membrane surface by the hydrophobic effect of cholesterol. Subsequently, $1\times10^5$ CEM cells were added and incubated for another 30 min at 37 °C with a metal bath shaking at 300 rpm. The nanobrush is again anchored to the CEM cell membrane surface by the hydrophobic effect of cholesterol, and finally, the cell assembly is obtained. The NIKON A+ confocal microscope observed cell assembly with 490 nm laser and 540 nm laser excitation. The cell images were observed with 100 × objective lens. Quantitative data of cell assembly were derived using flow cytometry with 490 nm channel and 540 nm channel excitation. The above operation was repeated 3 times.

##### 1.6. Twisted DNA nanobrush for prolonged incubation to build cell clusters

Individual CEM cells to form cell clusters: $1\times10^6$ CEM cells were mixed in 5 mL of 1640 medium, placed in a culture bottle and cultured overnight for 12 h. Among them, 1 mL of CEM cells was taken and centrifuged at 1200 rpm for 3 min. We resuspended the cell in 200 μL of 1640 medium and adjusted the cells to $1\times10^5$ for subsequent use. Then CEM cells were added nanobrush (b3-2chol) at a final concentration of 500 nM and placed in the incubator at different times. CEM cells will form stable cell clusters within 24 hours under the combined effect of hydrophobic insertion. Photographs were taken at different time points using a cell microscope under bright field conditions. The magnification of the microscope was 10×.

**1.7. Working principle of MIET and lifetime-distance conversion**

The principle of MIET has been elaborated in our previous publications (ref 1-5). Similar to the fluorescence resonance energy transfer (FRET) process, the gold film can work as an efficient energy acceptor of the excited state energy of a fluorophore. This leads to a strongly distance-dependent modulation of lifetime of the fluorophore over a distance range of ~ 150 nm above the gold surface. This modulation can be calculated by modeling the emitting fluorescent molecule as an electric dipole emitter and solving Maxwell's equation with this source field in the presence of the MIET substrate. Taking into account also the non-radiative transition rate, the observable excited-state fluorescence lifetime ($\tau_f$) is then found as

$$\frac{\tau_f(\theta, z_0)}{\tau_0} = \frac{S_0}{\phi S(\theta, z_0) + (1-\phi)S_0}$$

where $S(\theta, z_0)$ is the emission power of the dipole emitter at the distance of $z_0$ from the substrate surface with orientation angle of $\theta$, $\tau_0$ is the free-space fluorescence lifetime in absence of the gold film, $\varphi$ represents the quantum yield of the fluorophore, and $S_0$ is the free-space emission power of an ideal electric dipole emitter given $S_0 = cnk_0{}^4p^2/3$ with c being the speed of light, $k_0$ the wave vector amplitude in vacuum, n the refractive index of water, and p the amplitude of emission dipole moment vector. MIET exploits this lifetime-to distance ($\tau_f$ versus z) dependence for converting measured lifetime values into distance values, see Figure 3 in main text. To calculate this model curve, *a priori* knowledge of the fluorophore's $\varphi$, $\tau_0$, and $\theta$ is required. Previously, we have determined these values for DPPE-atto655: $\varphi = 0.36$, $\tau_0 = 2.6$ ns.

The fluorophore orientation we used for the GUVs measurement is parallel to the membrane and for the cell measurement we used a random orientation.

For the conversion of fluorescence lifetimes into distance values, we used the calculated lifetime-versus-distance MIET curve as described above (see also Figure 3 in main text). For this purpose, a custom-written MATLAB script was used. A MATLAB-based software package for the calculation of MIET lifetime-versus-distance curves as well as the conversion of a lifetime to distance, equipped with a graphical user interface, has been published (ref 6) and is available free of charge at https://projects.gwdg.de/projects/miet. While the published version of the software assumes that the fluorescent emitters are rotating quickly compared to their excited-state lifetime, this was not the case for the measurements in the present work. Here, a dye orientation parallel to the bilayer (and thus to the substrate) was assumed when calculating the MIET calibration curve.

##### 1.8. Intermembrane distance measurement of model membrane

For MIET measurement, 10 μL of 1 μM nanobrushes were added to the SLBs and incubated for 30 min, followed washing by copious buffer to remove the unbonded nanostructures. The diluted GUVs were added onto the nanostructures and incubated for 30 min.

##### 1.9. Real-time observation of cell surface engineering process by MIET

Mouse embryonic fibroblast cells (NIH-3T3 cell) were cultured in DMEM medium (GIBCO), which was supplemented with 10% fetal bovine serum (FBS) and 1% antibiotics (penicillin-streptomycin-amphotericin B) at 37 °C in a $CO_2$ incubator.

For real-time observation of cells, similar to the GUV measurement, SUVs were deposited on the MIET substrate and fused at room temperature for 30 min to form a uniform lipid bilayer. The substrate was then washed with copious PBS buffer to remove unbound vesicles. Next, 10 μL of 1 μM nanobrush was added to the SLBs and incubated for 30 minutes. After incubation, the substrate was washed with PBS buffer to remove the unbounded nanobrush. Liposome-stained NIH-3T3 cells were then added to the system. Once the cell settled onto the SLB, continuous scanning of the sample was started. A stage top incubator (Cat. No: 12722, Silver Line, ibidi) fits in the microscope, connects to incubator temperature, gas and humidity controllers and creates the proper environment (37 °C, 5% $CO_2$) for live-cell imaging right on the microscope stage.

#### 2. Schematic diagram of DNA nanobrush

(A) DNA nanobrush with backbone 21 bp (b1)

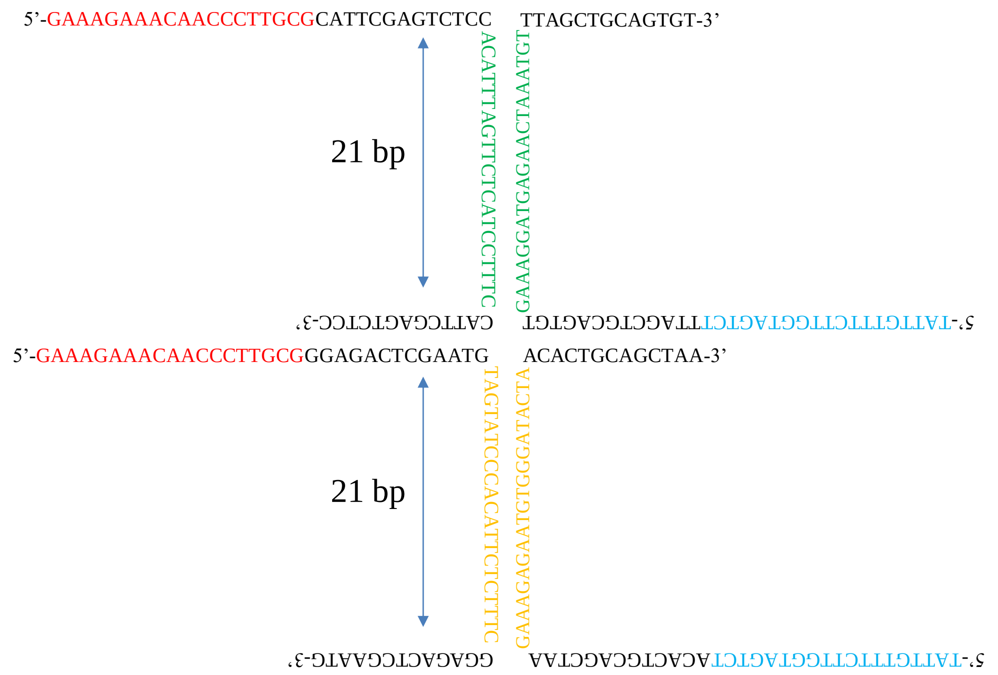


(B) DNA nanobrush with backbone 21 bp ligated cholesterol (b1-2chol)

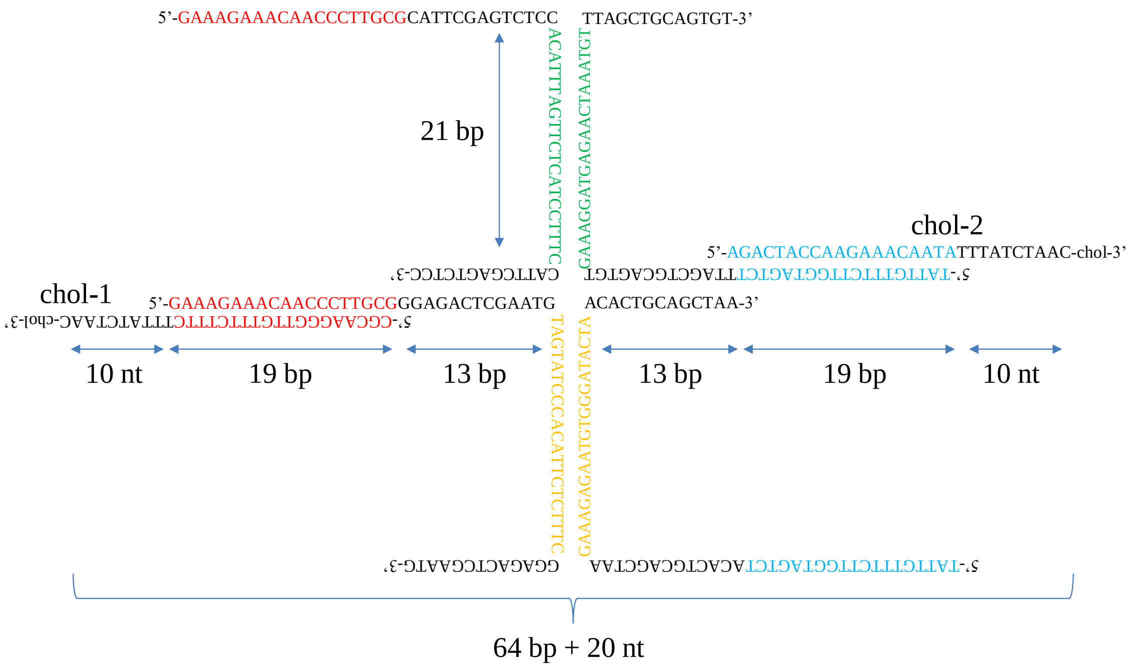

(C) DNA nanobrush with backbone 22 bp ligated cholesterol (b2-2chol)

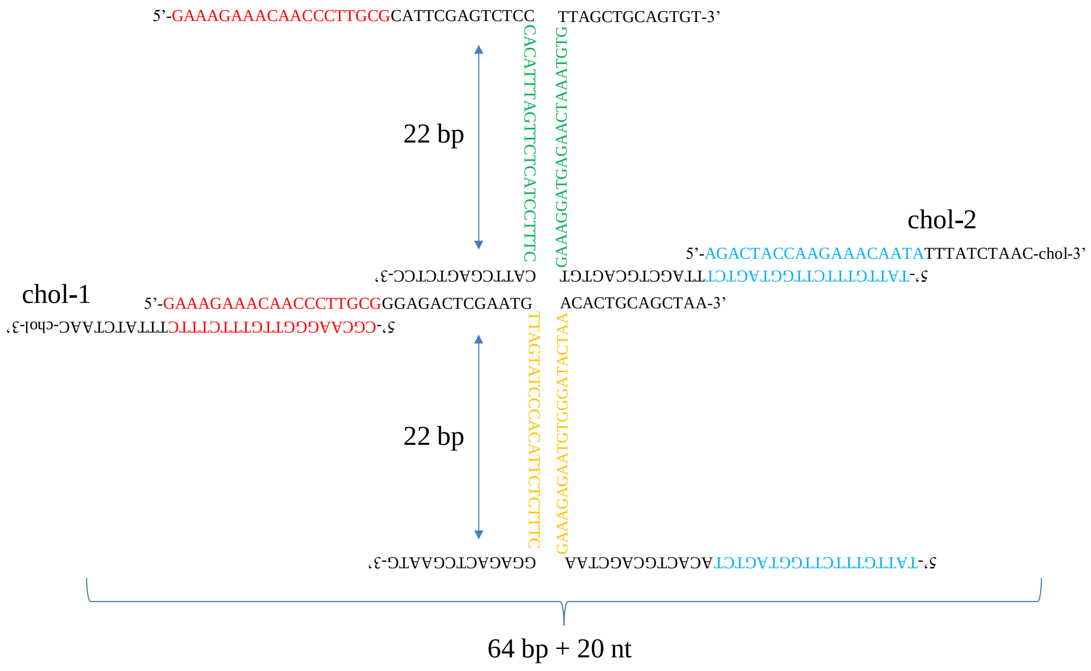


(D) DNA nanobrush with backbone 25 bp ligated cholesterol (b3-2chol)

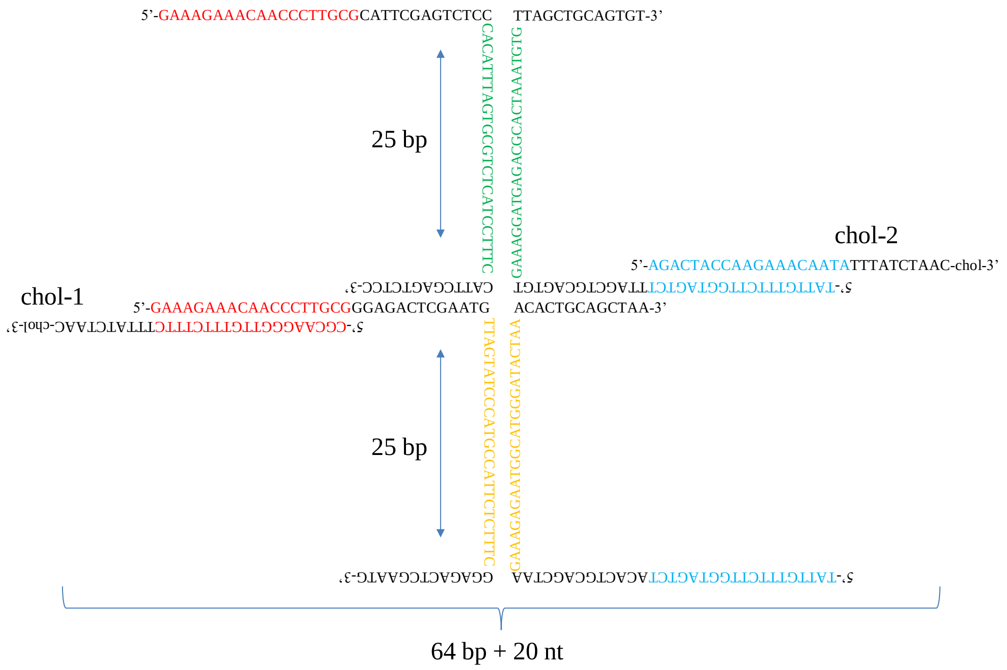

(E) Short side arm DNA nanobrush with backbone 21 bp ligated cholesterol (b4-2chol)

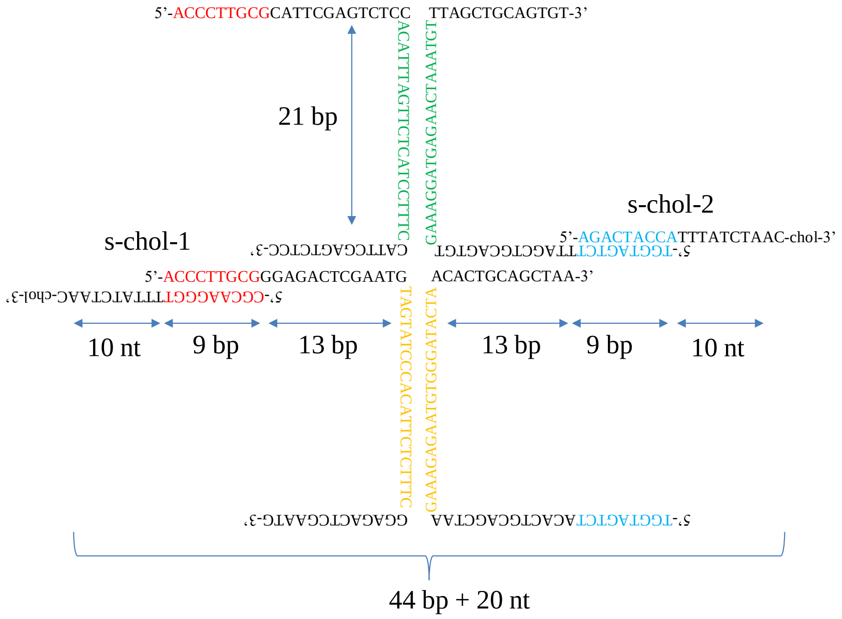


(F) DNA nanobrush with backbone 22 bp ligated one side cholesterol (b2-1chol)

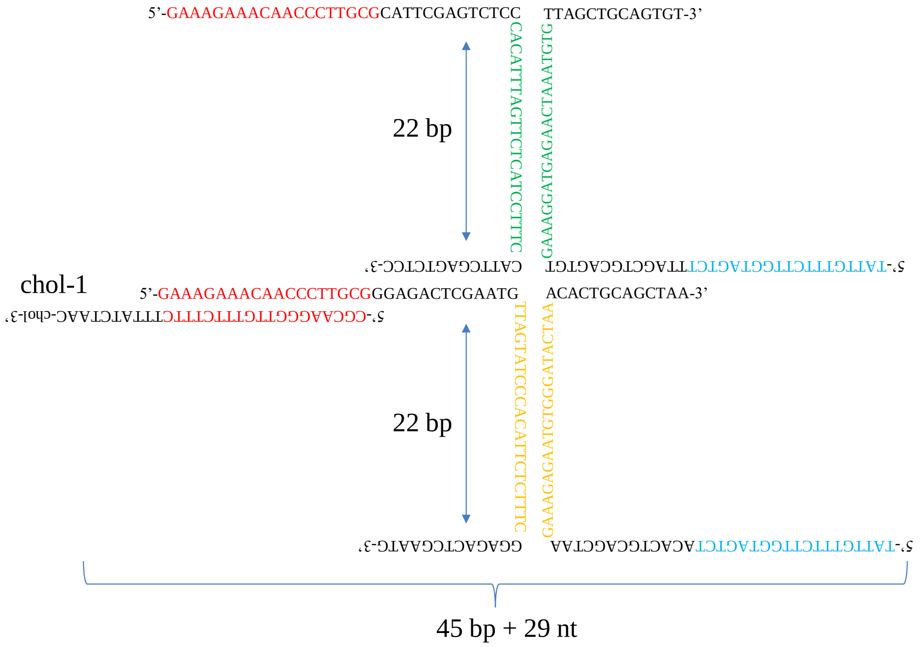


**Figure S1.** Schematic diagram of DNA nanobrush. (A) The nanobrush with a backbone 21 bp is a planar structure (b1). (B-D) Changing the backbone bases of the nanobrush to regulate the recognition direction. The nanobrush with a backbone 21 bp is a planar structure. The nanobrush with a backbone 25 bp is a completely twisted conformation. (E) Nanobrush with the short arm where the backbone is 21 bp (b4), both side arm connects cholesterol (b4-2chol). (F) Nanobrush with backbone 22 bp, one side arm connects cholesterol (b2-1chol).

#### 3. Synthesis and characterization of the DNA nanobrush

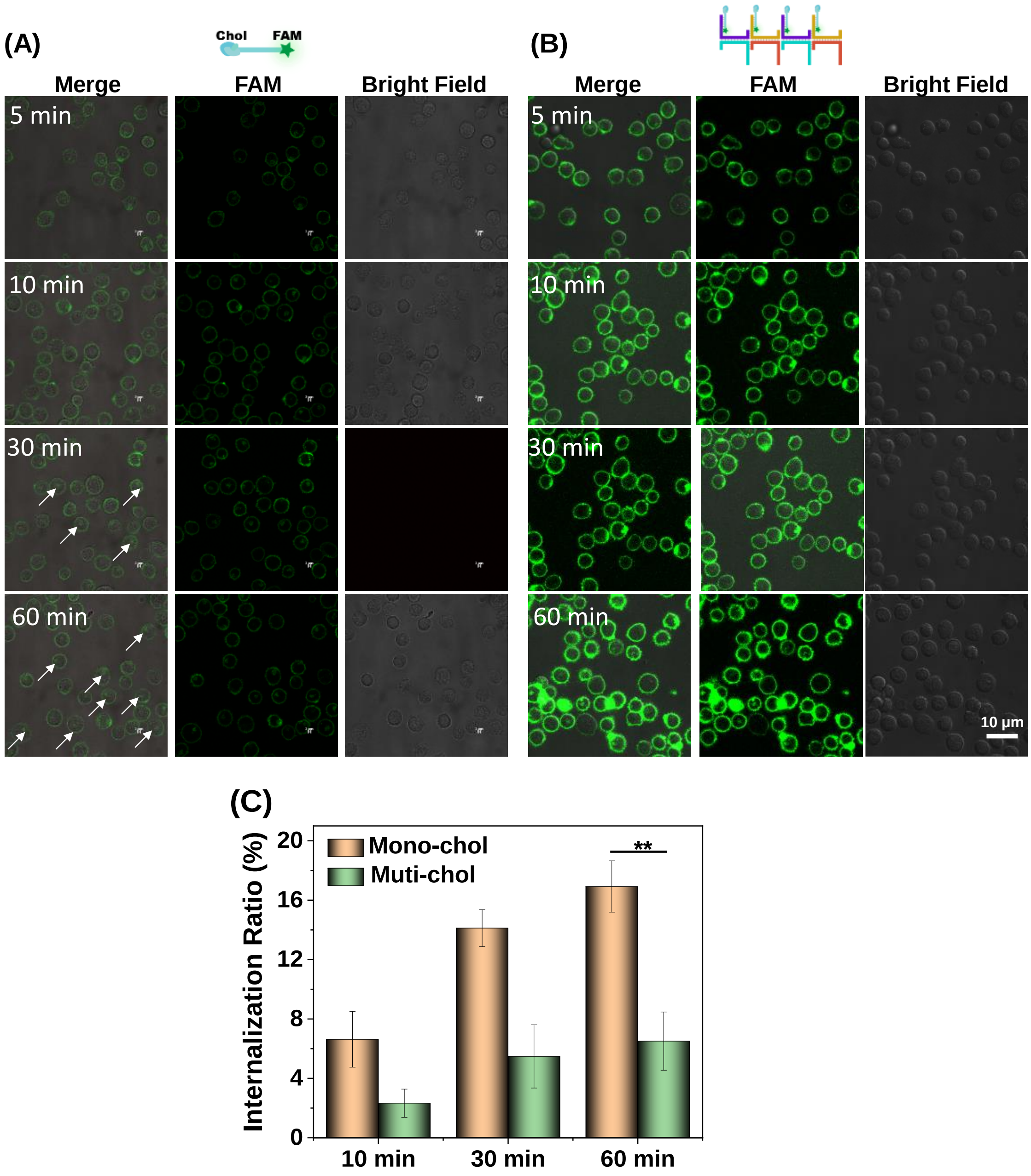


**Figure S2.** Internalization of nanobrush under long-time incubation. Confocal laser scanning microscopy (CLSM) images showed the internalization of monovalent (A) and multivalent (B) cholesterol. Reactions were performed at 37 °C for different times in a medium containing 10% FBS. (C) Relative cell internalization rate analysis was performed on the corresponding samples. **P ≤ 0.01 by unpaired two-tailed t-test.

For the statistical analysis, over 100 cells were measured with Image J software for each sample group, and at least three independent experiments were performed.

#### 4. Optimization of two kinds of cells assembled by nanobrush

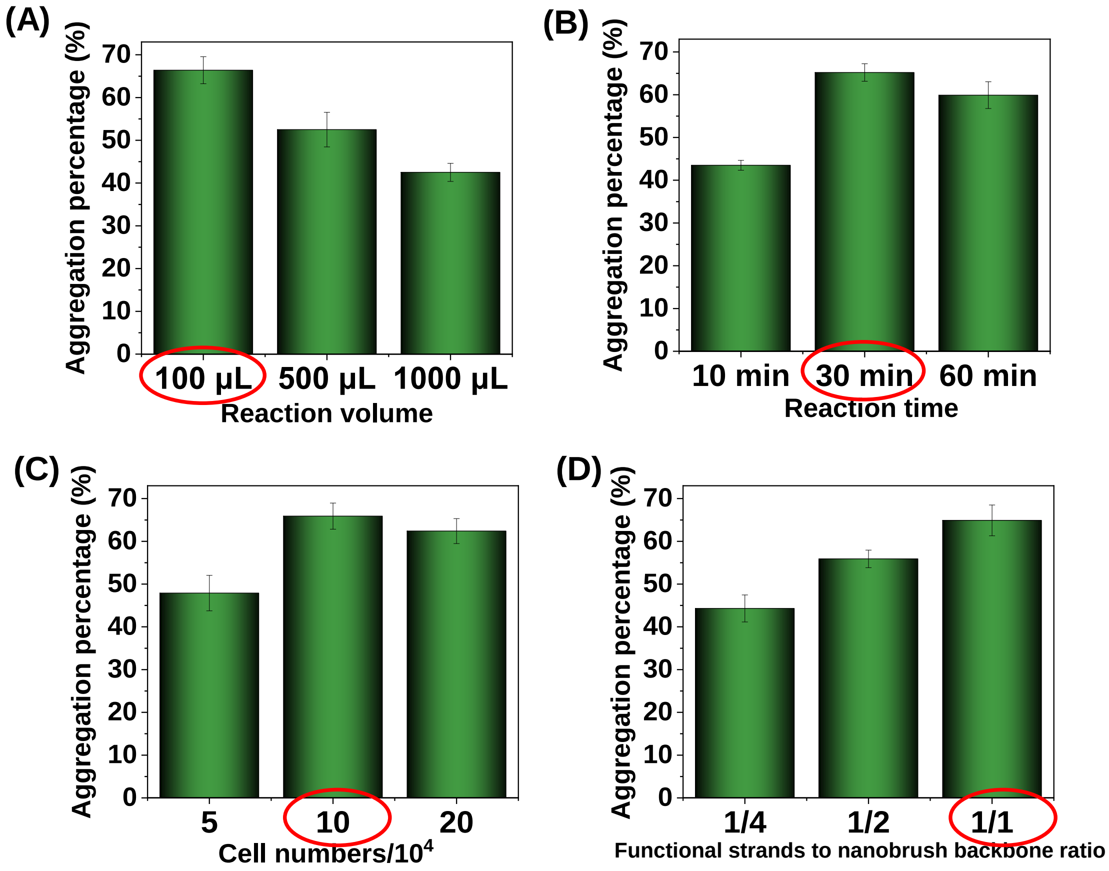


**Figure S3.** Optimizing the reaction conditions during two kinds of cell assembly: reaction volume (A); reaction time (B); cell number (C); the proportion of functional strands to nanobrush backbone sites. The optimal reaction conditions were: $1 \times 10^5$ cells were reacted in 100 μL for 30 min. Also, the assembly efficiency is maximized when the functional strand completely occupies each side arm of the backbone extension. Bars represent mean ± SD ($n = 3$).

#### 5. Scheme of MIET experimental set-up

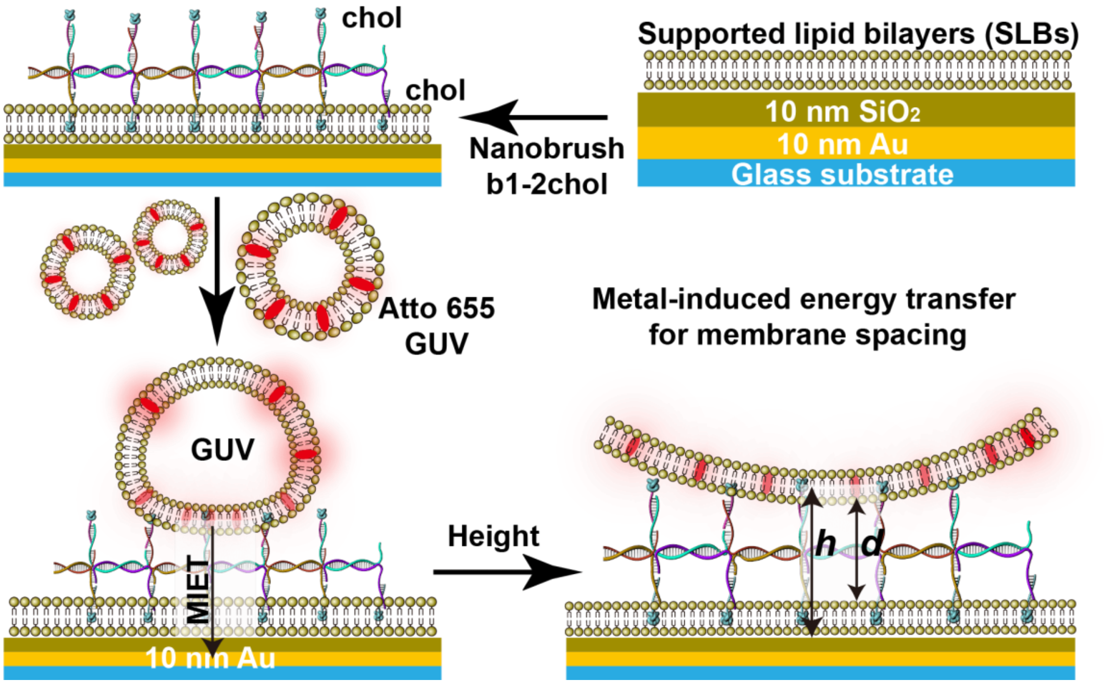


**Figure S4.** Scheme of the experimental set-up. The SLBs and GUVs were used to detect the distance between the two membranes assembled by the nanobrush.

#### 6. MIET measurements of DNA nanobrush-assembled membranes

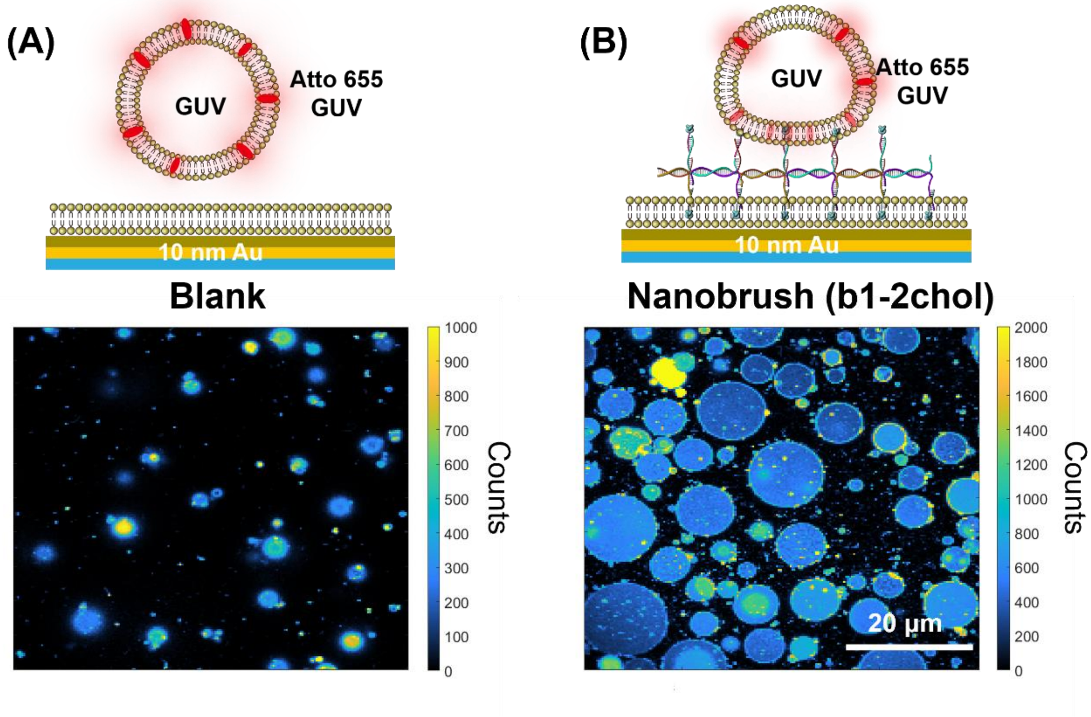


**Figure S5.** Fluorescence intensity images showed the adherence of the GUVs membrane without (A) or with (B) the addition of nanobrush. At least three independent experiments were performed.

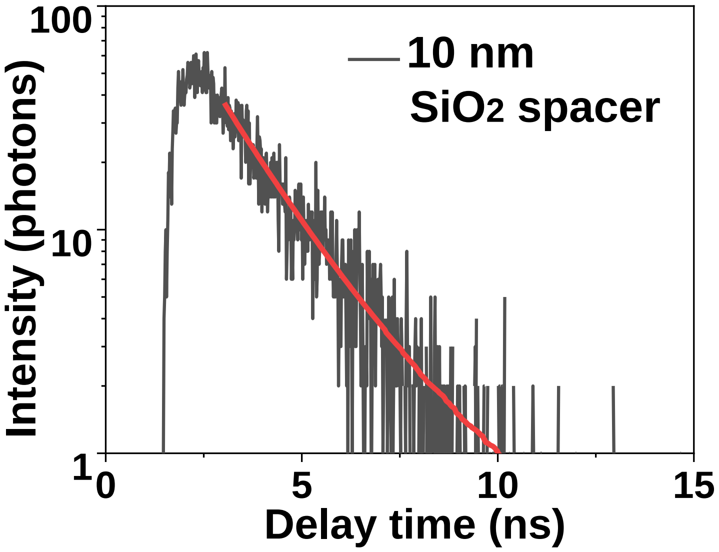


**Figure S6.** Typical time-correlated single photon counting (TCSPC) histogram of a single pixel (black). Multi-exponentially fit (red) to obtain the fluorescence lifetime.

#### 7. MIET microscopy to visualize the binding process using NIH-3T3 cells

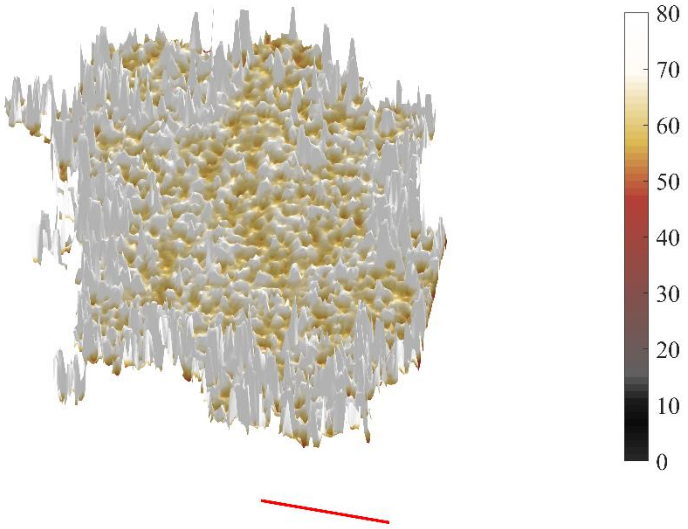


**Figure S7.** Three-dimensional height reconstruction of NIH-3T3 cells at 60 min in the absence of nanobrush. The scale bar is 5 µm.
